# Molecular Complexes for Effective Inhibition of Tau Aggregation

**DOI:** 10.1101/363572

**Authors:** Nalini V. Gorantla, Vinod G. Landge, Pramod G. Nagaraju, Lisni P. Sunny, Anjhu Nair, Siba P. Midya, Poornima Priyadarshini CG, Ekambaram Balaraman, Subashchandrabose Chinnathambi

## Abstract

Tau is an axonal protein known to form abnormal aggregates and is the biomarker of Alzheimer’s disease. Metal-based therapeutics for inhibition of Tau aggregation is limited and rarely reported in the contemporary science. Here, the first example is reported of a rationally designed molecular cobalt(II)-complexes for effective inhibition of Tau and disaggregation of preformed Tau fibrils. The mechanistic studies revealed that the prevention of Tau aggregation by CBMCs is concentration-dependent and Tau seldom exhibits conformational changes. Interestingly, CBMCs play a dual role by causing disassembly of preformed aggregates as well as complete Tau inhibition. We believe that this unprecedented finding by the newly developed molecular complexes has a potential to lead to developing innovative metal-based therapeutics for Alzheimer’s disease.

## INTRODUCTION

Alzheimer’s disease (AD) is a neurodegenerative disorder characterised by progressive cognitive and behavioural impairment. Worldwide 44 million people are known to have AD and its related dementia. Abnormal protein deposits in the brain, such as extracellular amyloid plaques and intracellular neurofibrillary tangles (NFTs), characterize AD. The microtubule-associated protein Tau (MAPT) plays a key role in several neurodegenerative diseases, like AD, FTDP-17 and Parkinson’s disease *etc*.^1^ The axonal protein Tau is expressed in the adult human brain as six different isoforms. Due to alternative splicing, two N-terminal inserts and the second repeat (R2) in the C-terminal microtubule-binding domain can be present or absent (Fig. 1A). Upon hyperphosphorylation, Tau disassembles from microtubule (MT) and self-assembles to form NFTs which consist of straight paired helical filaments (PHFs).^1^ Several factors are responsible for triggering under pathological conditions, such as post-translational modifications, oxidative stress, truncation, and metal ions *etc*.^3,4^ Thus, there is an immense interest to identify potentially active molecules derived from metal complexes, natural products, and short-range peptides for inhibition of Tau aggregation or to disaggregate the pre-formed fibrils of Tau.^5,6^ Recently, methylene blue and methylthionine hydrochloride have been identified as inhibitors of Tau aggregation and were subjected to phase III clinical trials.^7^ Indeed, there is a strong need for the discovery of new potential therapeutics.^8-9^ In this regard, significant research is being devoted in recent times to identify active compounds^10,11,12,13^ with novel scaffolds that may have potential properties for the treatment of AD by inhibiting Tau aggregation.^14,15^

**Figure 1.**
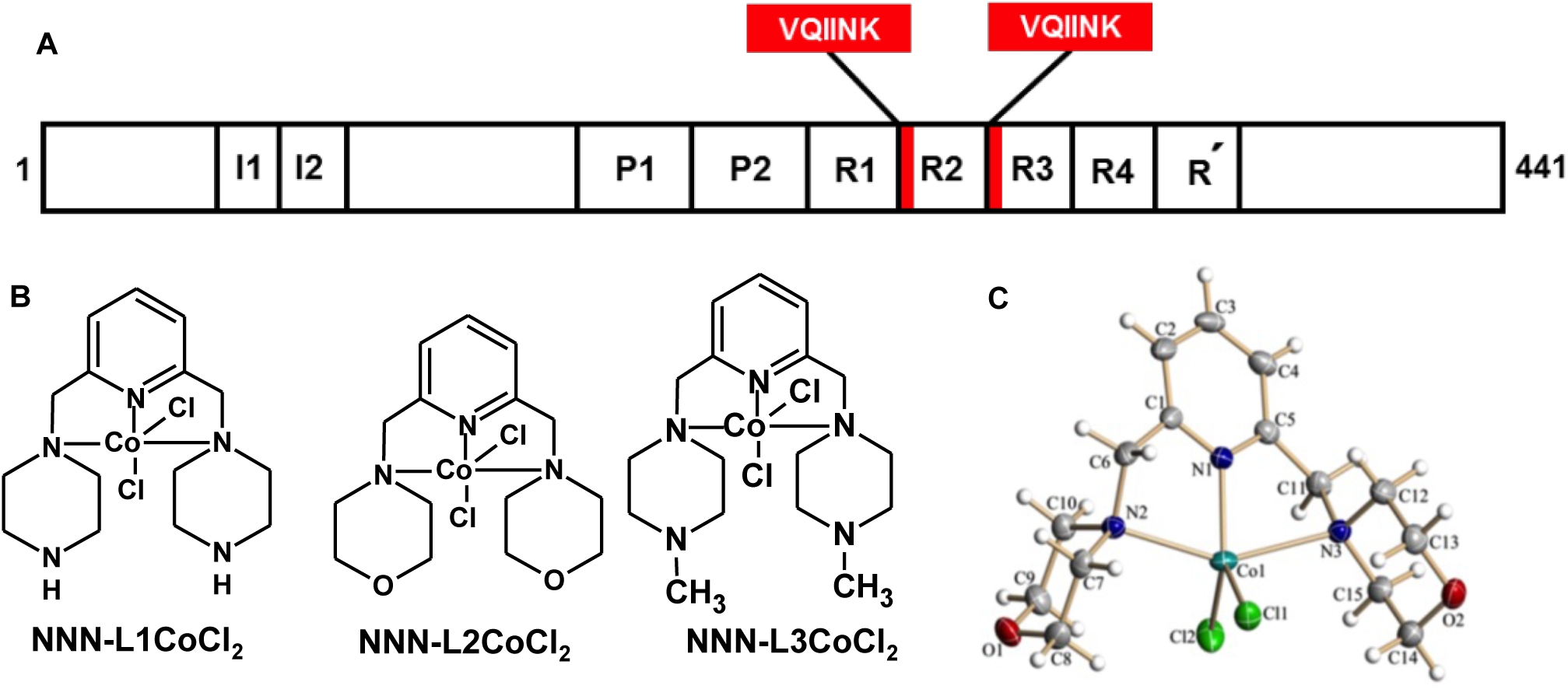
(a) Domain organization of full-length Tau. The longest isoform of Tau is composed of 441 amino acids, with four repeats towards C-terminal that is crucial in both physiological processes as well as in AD pathology. In physiological conditions, it interacts with the tubulin dimer and helps in the assembly of MTs, while in the pathological condition they act as main nucleating centers and form the core of aggregates. The four repeats, R1 to R4 comprises of hexapeptides at the beginning of R2 and R3 which are the signature motifs responsible for Tau aggregation. (b) Chemical structure of cobalt-based metal complexes. (c) X-ray crystal-structure analysis of NNN-L2 CoCl_2_ with 50% probability of thermal ellipsoids. Selected bond length [A°] and angle [°]: Co(1)-N(1) 2.0265, Co(1)-N(2) 2.3215, Co(1)-N(3) 2.4530, N(1)-Co(1)-N(2) 76.97, N(1)-Co(1)-N(3) 73.51, Cl(1)-Co(1)-Cl(2) 113.56.

Recently, the importance of different metals in the AD is well studied, which elucidates the critical role of metal ions in pathology.^16^ Metal ions are mostly involved in the physiological functions and they are also known to interact with proteins leading to their aggregation.^17,18^ Metal ions such as copper(II), zinc(II), iron(III) and aluminium(III) are well-studied for their protein aggregation property. It was reported that copper and zinc interact with Amyloid β and promote their aggregation.^19,20^ The mode of binding of these metal complexes with Amyloid β was studied by Raman spectroscopy and revealed the importance of the three histidines present at the N-terminus.^21^ Later, the role of iron(III) in the aggregation of Amyloid β was studied by NMR experiments, which revealed the importance of first 16 amino acids in the formation of iron coordination.^22^ The interaction of copper(II) ions with Tau was analyzed by Soragni *et al*., by NMR studies, which showed the importance of amino acid sequences adjacent to hexapeptide motif in the second and third repeat of Tau. The effect of copper on Tau protein aggregation *via*. oxidation of cysteine residues was the key interest for copper binding.^23^ In AD brain, Tau hyperphosphorylates and aggregates to form PHFs, ions such as aluminium(III) specifically interact with its phosphorylated epitopes.^24^ These findings suggest that phosphorylated Tau is triggered for aggregation in the presence of aluminium(III). Further, the effect of aluminium maltolate administration was reported for neurodegeneration in rabbits.^25^ However, the effect of zinc(II) on Tau fibrillization in physiological conditions is contradicting to that of copper(II) and aluminium(III). The lower levels or deficiency of zinc(II) would enhance Tau aggregation by decreasing the expression of Tau or by reducing the ability of Tau to interact with MTs. Sequestration of zinc(II) by extracellular senile plaques leads to decreased intracellular zinc levels, which ultimately leads to NFTs formation. Importantly, the presence of higher intracellular zinc levels leads to hyperphosphorylation of Tau at S214, which eventually forms aggregates.^26^ Furthermore, ferric iron can lead to Tau aggregation *in vitro*, and its reduced form causes hyperphosphorylation of Tau.^27,28^ Other targets of AD such as acetylcholine esterase and monoamine oxidase are also altered by the levels of metal ions.^29^ Notably, other neurodegenerative diseases like Creutzfeldt Jakob disease are also known to be caused by the presence of unbalanced metal ions. The cellular prion protein PrP^c^ undergoes conformational changes to PrP^Sc^ and leads to prion disease. Similarly, the pathological conformational changes in the prion protein were observed upon its interaction with copper(II).^30^ The significance of copper interaction with octarepeats in PrP was revealed by using NMR, tryptophan fluorescence emission and Raman spectroscopy.^30,31^

In our present studies, we screened the effect of rationally designed molecular cobalt based metal complexes (CBMCs; Fig. 1B) on Tau aggregation and found that these complexes are effective in preventing the formation of a toxic population of Tau. The CBMCs were effective in inhibiting polymerization of Tau in concentration-dependent manner. The efficacy of these metal complexes were analysed by fluorescence assay, studying the conformational change in Tau by spectroscopic techniques and the morphology of aggregates was observed by microscopic analysis.

## RESULTS AND DISCUSSION

### CBMCs, as a barrier for Tau aggregation

Tau protein is one of the major microtubule-associated proteins in neuronal axons that mainly functions to stabilize and assemble MTs.^32^ Tau is a soluble protein and adopts natively unfolded structure in solution.^33^ The repeat domains of Tau and the flanking proline-rich regions confers the property of MT binding and assembly (Fig. 1A). The repeat domains of Tau represents the core of PHFs.^34^ The hexapeptide motifs in repeat 2 and 3 form the basic motif for aggregation. *In vitro* conditions, heparin is used as an inducer to enhance Tau aggregation.^34^ Heparin binds to the positively charged residues of flanking repeats 2 and 3, and thus leading to charge neutralization. Further, the binding of heparin also leads to change in conformation of β-sheet, which serves as a nucleation centre for the aggregation. Tau protein was diluted in assembly buffer, incubated for the formation of aggregates in presence or absence of CBMCs and the extent of aggregates formation was monitored by ThS. Our interest in metal-based therapeutics in AD began with the discovery of novel Co(II) complexes, since cobalt is relatively abundant, and biorelevant. The reaction of the tridentate ligand with CoCl_2_ in methanol at 65 °C for 4 hours results the corresponding cobalt-pincer complexes in excellent yields (See ESI, S1-S7). All the complexes were well-characterized and the structure of NNN-L2 CoCl_2_ was confirmed by a single-crystal X-ray diffraction study (Fig. 1C).

The Co(II)-complexes (CBMCs) were incubated with constant Tau concentration of 0.91 mg mL^−1^, with increasing concentrations of metal complexes (0.01, 0.025, 0.05 and 0.1 mg mL^−1^). The aggregation of full-length Tau with CBMCs at the higher concentration substantially decreased the ThS fluorescence (Fig. 2A, B and C), which indicates the prevention of aggregates formation. Further, we have quantified inhibition, which revealed that L2 is more potent with 92.5% inhibition at the lowest concentrations of 0.025 mg mL^−1^, whereas, L3 and L1 showed 85.5 and 73.9% inhibition, respectively. At the highest concentration of 0.1 mg mL^−1^, they showed up to 93% inhibition (Fig. 2D). The higher order species formed upon Tau aggregation were observed by SDS-PAGE analysis (Fig. 2E, F and G). In presence of metal complexes, Tau was able to form higher order aggregates which completely faded away over the time. At 0 hour the intensity of Tau was high which decreased with increasing complex concentration (Fig. 2H). This data is in agreement with the size-exclusion chromatography analysis which is evidenced by the retention volume that metal complexes prevent the aggregation of Tau to higher molecular weight species *in vitro* (Fig. S12A, B, C and D).

**Figure 2.**
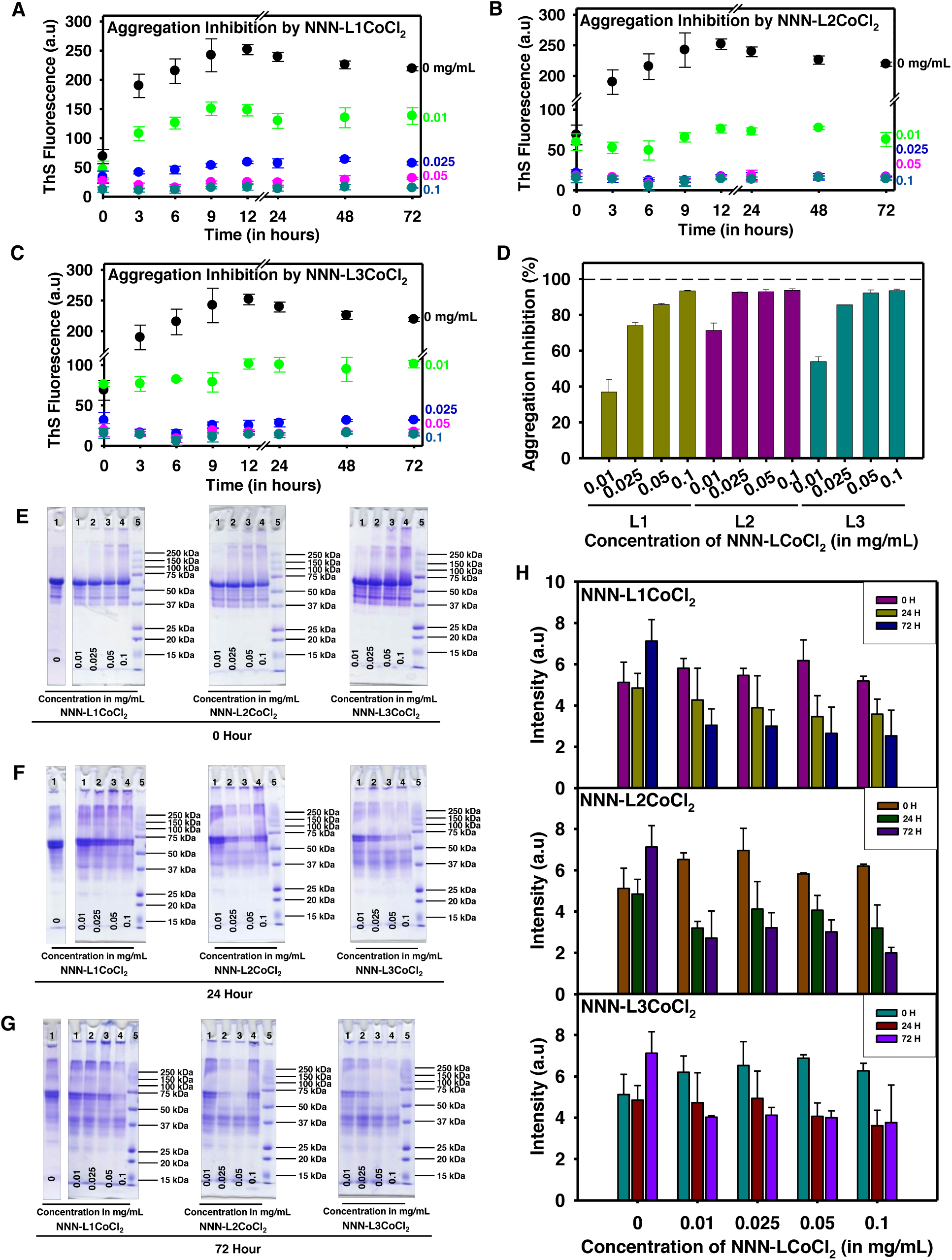
Cobalt-based complexes inhibits Tau aggregation. (a, b, c) The aggregation inhibition of full-length Tau in presence of CBMCs monitored by ThS fluorescence. The aggregation was induced from full-length Tau in presence of heparin as inducer and assembly buffer. The typical full-length Tau reaches its highest propensity of aggregation, but the incorporation of cobalt complex completely decrease its aggregation with increasing concentration of metal complex. (d) Although all three complexes are compelling in inhibiting aggregation process, the lower concentration of L2 is more effective in comparison with L1 and L3. At higher concentrations of 0.05 and 0.1 mg mL^−1^ L2 and L3 show maximum inhibition of about 97%. L1 at its highest concentration of 0.1 mg mL^−1^ also has an inhibition of 97%, which clearly suggests the proficiency of these metal complexes in inhibiting Tau aggregation. (e) Tau was analysed by SDS-PAGE at 0 hour, where the higher order species were observed in the highest concentration of CBMCs. (f) The higher order species were reduced at 24 hours. (g) When Tau was further analyzed at 72 hours these higher order species completely faded away. (h) The quantitative analysis of Tau in presence of CBMCs indicate the reduction in the formation of higher order aggregates with prolonged incubation (up to 72 hours), which reveals the potency of the complexes in preventing their aggregates formation.

### Native conformation of Tau maintained by CBMCs

The changes in Tau conformation due to metal complexes were studied by CD spectroscopy. Tau has no secondary structure and exists as a random coil protein, which absorbs around 190 nm. The aggregated Tau shows a shift towards the higher wavelength, which signifies the transition of the random coil to β-sheet conformation. In the presence of CBMCs, the typical random coil conformation of full-length (Fig. 3A, B and C) was not altered. These results signify the effect of these complexes in preventing aggregates formation. Tau protein *in vitro* forms fibrillar morphology in the presence of heparin, which is observed as a long filamentous structure. The extent of aggregate formation by Tau in presence of L1, L2 and L3 were studied by Transmission electron microscopy (TEM), which discloses the vigour of the CBMCs in completely preventing the aggregates formation (Fig. 3D, E, F and G). Further, the EDAX (Energy Dispersive X-Ray) revealed that the negligible amount of CBMCs was associated with Tau, which denotes that Tau was devoid of conjugation with CBMCs (data not shown). X-ray photon electron spectroscopy analysis showed that the oxidation state of the CBMCs remains unchanged during Tau aggregation (data not shown). The formation of higher order species by full-length Tau at 0 hour in presence of CBMCs stimulated us to study its role on soluble Tau. However, surprisingly no higher order aggregates were observed upon incubation at 37 °C (Fig. S9A) and this manifests the inability of the complexes to form toxic Tau species when compared with the control (Fig. S9B). The initial Tau conformation plays a critical role in order to form the nucleus, which further accelerates the formation of aggregates. Here, we show that these molecular Co(II)-complexes did not induce the conformational changes in soluble Tau and maintained its native conformation (Fig. S10A, B and C). To support the data, we further studied the effect of CBMCs on soluble full-length Tau (Fig. S11A, B and C) for their aggregation. Tau aggregates were not observed, which indicated that the Co(II)-complexes did not lead to the formation of aggregates during initial stages of Tau assembly.

**Figure 3.**
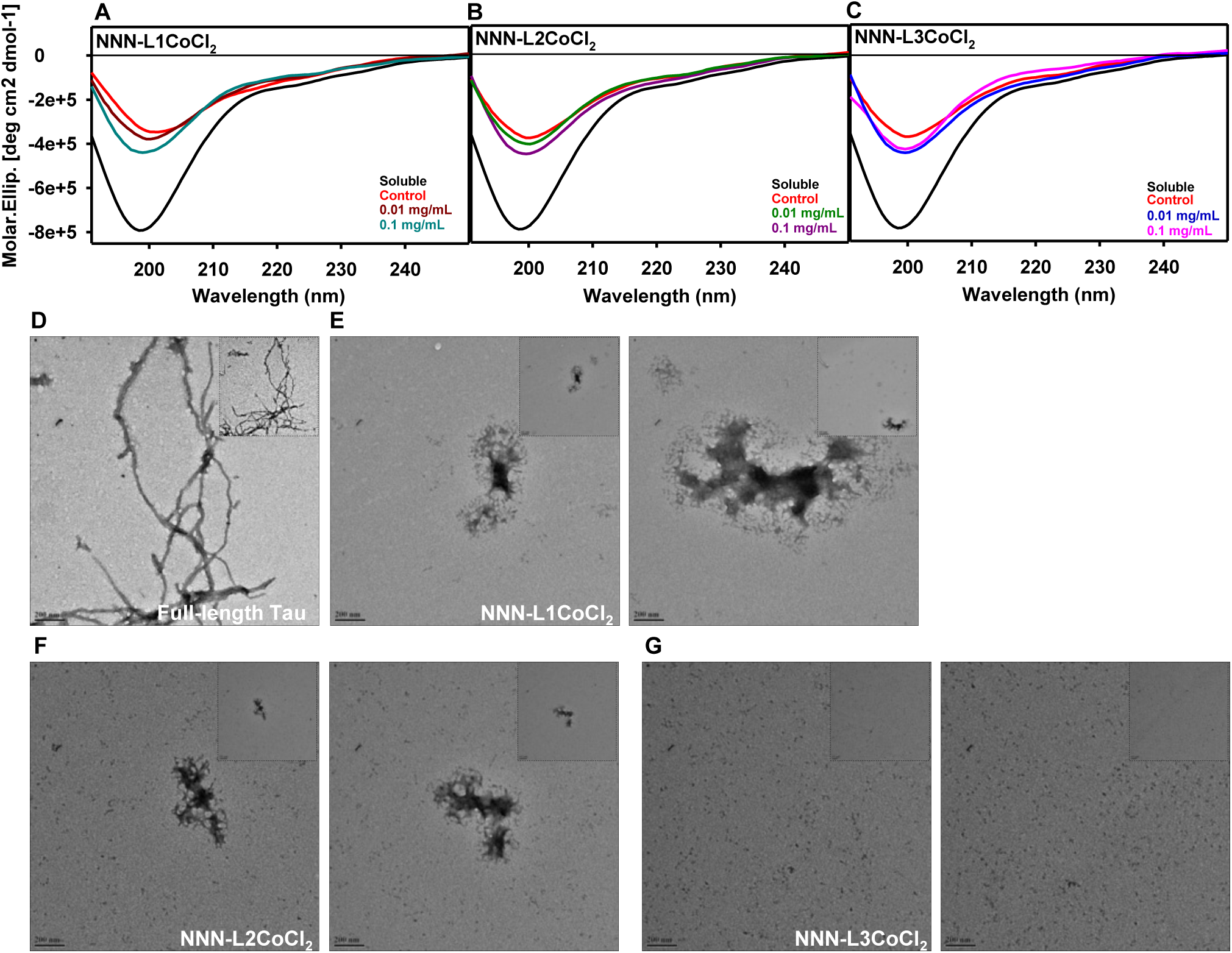
Conformation of full-length Tau measured by CD spectroscopy. (a, b, c) The full-length Tau in native state unveils its random coil conformation, but upon aggregation it attains β-sheet conformation, represented in red. On addition of CBMCs in increasing concentrations of 0.01 and 0.1 mg mL^−1^ Tau exhibits random coil conformation, indicating that CBMCs are effective in preventing β-sheet formation by Tau and thus, preventing its aggregation. (d) The morphology of Tau upon incubation with inducer alone and the typical morphology of Tau fibrils were observed. (e, f, g) The presence of CBMCs stipulated their ability to prevent aggregate formation. The inserts in each micrograph represent the morphology of Tau aggregates at a magnification of 0.5 μm.

### Destabilization of preformed Tau fibrils

We further investigated the role of CBMCs on disassembly of Tau PHFs. This would enhance both properties of aggregation inhibition and disaggregation of Tau. The preformed Tau aggregates were incubated with various concentrations of L1, L2 and L3 and it was observed that L3 was more effective in disaggregating Tau when compared to L1 and L2 (Fig. 4A, B and C). At a concentration of 0.5 mg mL^−1^, L3 showed about 77.4% of inhibition, but L1 and L2 showed 73.5% and 71.9% inhibition, respectively (Fig. 4D). Furthermore, SDS-PAGE analysis showed a decrease in the aggregates by CBMCs in time dependent manner (Fig. 4E, F and G). Unlike the ThS fluorescence, SDS-PAGE did not show decrease in the intensity of higher order aggregates of Tau. However, at a concentration of 0.5 mg mL^−1^ L3 exhibited decrease in Tau aggregates and at 24 hours the decrease in aggregates by L3 was prominently observed. At 120 hours of incubation, CBMCs effectively disintegrated Tau aggregates as ascertained by the lower intensity of higher molecular weight Tau around 250 kDa (indicated by red arrow). Initially, at 0 hour of incubation, no changes were observed in the morphology of aggregates in presence of CBMCs (Fig. S8B, C and D), when compared to untreated Tau aggregates (Fig. S8A). However, at the end of 120 hours, there was a definite decrease in the formation of PHFs (Fig. 6A, B, C and D). Overall, these results suggest the efficacy of CBMCs in preventing the formation of Tau aggregates that could be toxic and hence, it can be a potent therapeutic agent to AD.

**Figure 4.**
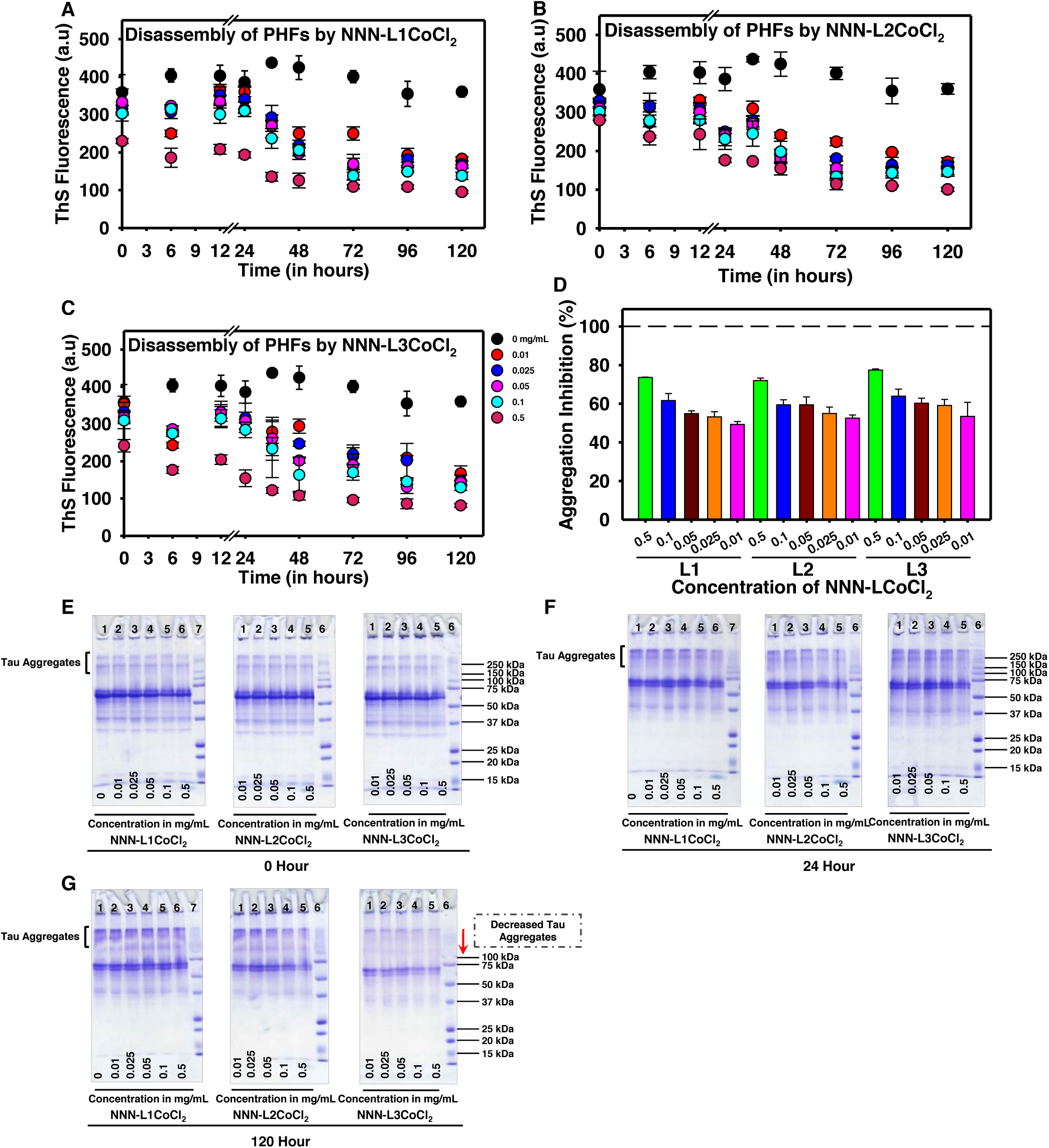
Disaggregation of full-length Tau. (a, b, c) The effect of disaggregation by L1, L2 and L3 conjugated to CoCl_2_ was analysed by ThS fluorescence assay. Tau aggregates were incubated with CBMCs in an increasing concentration of 0.01 mg mL^−1^ to 0.5 mg mL^−1^. (d) It was observed that the highest concentration of L1 showed 73.5% disaggregation, whereas L2 and L3 showed 71.9 and 77.4% disaggregation respectively. (e, f, g) The presence of higher order aggregates were monitored by SDS-PAGE at different time points of incubation, at 0 hour (e) 24 hours (f) and 120 hours (g) where L3 was more efficient amongst the three in disintegrating Tau aggregates.

In recent years, several drugs failed in the clinical trials which emphasize the urge to develop and screen different natural as well as synthetic molecules to target AD pathology. Although AD has multifaceted effectors like kinases, proteases, oxidative stress *etc*., the structure of the protein plays very important role in interaction and further change in its conformation can lead to either oligomers or aggregates formation. It has been suggested that the oligomeric precursors to amyloid fibrils may substantially be more toxic than the fibers themselves. Even if this was the case, amyloid fibrils are likely to play an important role, either as reservoirs or sinks to toxic oligomers. Hence, a detailed screening of compounds could help in discovering potential drugs that prevent protein aggregation in AD. The role of metal ions in association with proteins and leading to their aggregation is a well-known phenomenon.^35^ Metal ions such as copper(II), zinc(II), aluminium(III) and iron(III) are known to cause protein aggregation, but our results suggest that molecular cobalt(II) complexes have a paradoxical effect. Previously, Rajendran *et al*., have reported the deposition of metals such as iron, copper and zinc in the AD brain, which revealed increased metal accumulation in comparison with healthy brain tissue.^36^ Aluminium is known for the accumulation of Tau in cultured neurons and AD pathology.^37^ The animal studies revealed that injecting PHFs with and without aluminium salts led to distinct effects. The aluminium salts caused the deposition of Amyloid β and several other proteins along with Tau.^38^ Moreover, the *in vitro* studies also showed that aluminium caused the formation of higher order aggregates, as examined by SDS-PAGE. Aluminium was known to cause aggregation in phosphorylated Tau. Aluminium also effects Tau aggregation by reducing the activity of PP2A, an enzyme essential for dephosphorylation of Tau and activating kinases such as cdk5 and GSK3β.^39,40^ Kawahara *et al*., showed that prolonged exposure to aluminium leads to conformational changes in Amyloid β and its aggregation.^41^

In our present observations, the biochemical studies revealed the potency of CBMCs in preventing Tau polymerization. Copper interacts with both Tau and Amyloid β, driving the pathology of the AD. The oxidative property of copper that leads to Tau aggregation *via*. cysteine residues is well understood.^23^ Studies in 3XTg-AD mice showed that treatment with copper led to activation of cdk5/p25 which caused hyper phosphorylation of Tau.^42^ Voss *et al*., elucidated that copper led to hyper phosphorylation of Tau in SH-SY5Y cells, in Amyloid β independent manner.^43^ Copper is also known for reactive oxygen species (ROS) production in presence of Amyloid β which leads to cytotoxicity.^44 45^ Copper interacts with Amyloid β and leads to a reduction in the content of β-sheet conformation that would further form amorphous aggregates.^46^ In our current studies, CBMCs did not drive conformational changes in full-length Tau. The soluble Tau has also manifested the signature of typical random-coil conformation in presence of metal complexes, which indicates that CBMCs does not drive conformational changes in Tau. Tau usually accumulates to form filamentous aggregates, but aluminium leads to the formation of amorphous aggregates.^47^

In complement to the CD spectroscopy of soluble Tau, TEM analysis also suggests the absence of toxic Tau species. The interaction and affinity of the prion protein for copper and silver is dependent on their conformation of the protein.^48^ The SEC analysis clearly indicated that the higher order species of Tau aggregates was prevented by CBMCs (Fig. S12).

### Direct interaction of CBMCs with Tau

Isothermal titration calorimetry (ITC) is the direct method to obtain free energy (ΔG), enthalpy (ΔH) and entropy (ΔS) changes along with dissociation constant (K_D_) and number of binding sites (N) for ligand on the protein. Soragni *et al*., and Zhu *et al*., in individual studies employed ITC to analyse the binding of copper and lead, respectively.^23, 49^ The ITC titration yielded differential power values during interaction of Tau with L2 (Fig. 5A). These values were integrated and exothermic peaks were obtained (Fig. 5B). The negative ΔG value, −7.51 revealed the spontaneity of binding. The titration suggested binding of L2 to Tau with dissociation constant (K_D_) of 5.09 μM ± 5.84 μM, a weak interaction. The fitting by one set of sites resulted in the N value (number of sites on Tau) of 0.64 ± 0.132. The binding indicated small enthalpy changes (ΔH) with negative entropy of −0.654 ± 0.226 and −6.86 kcal/mol, respectively.

**Figure 5.**
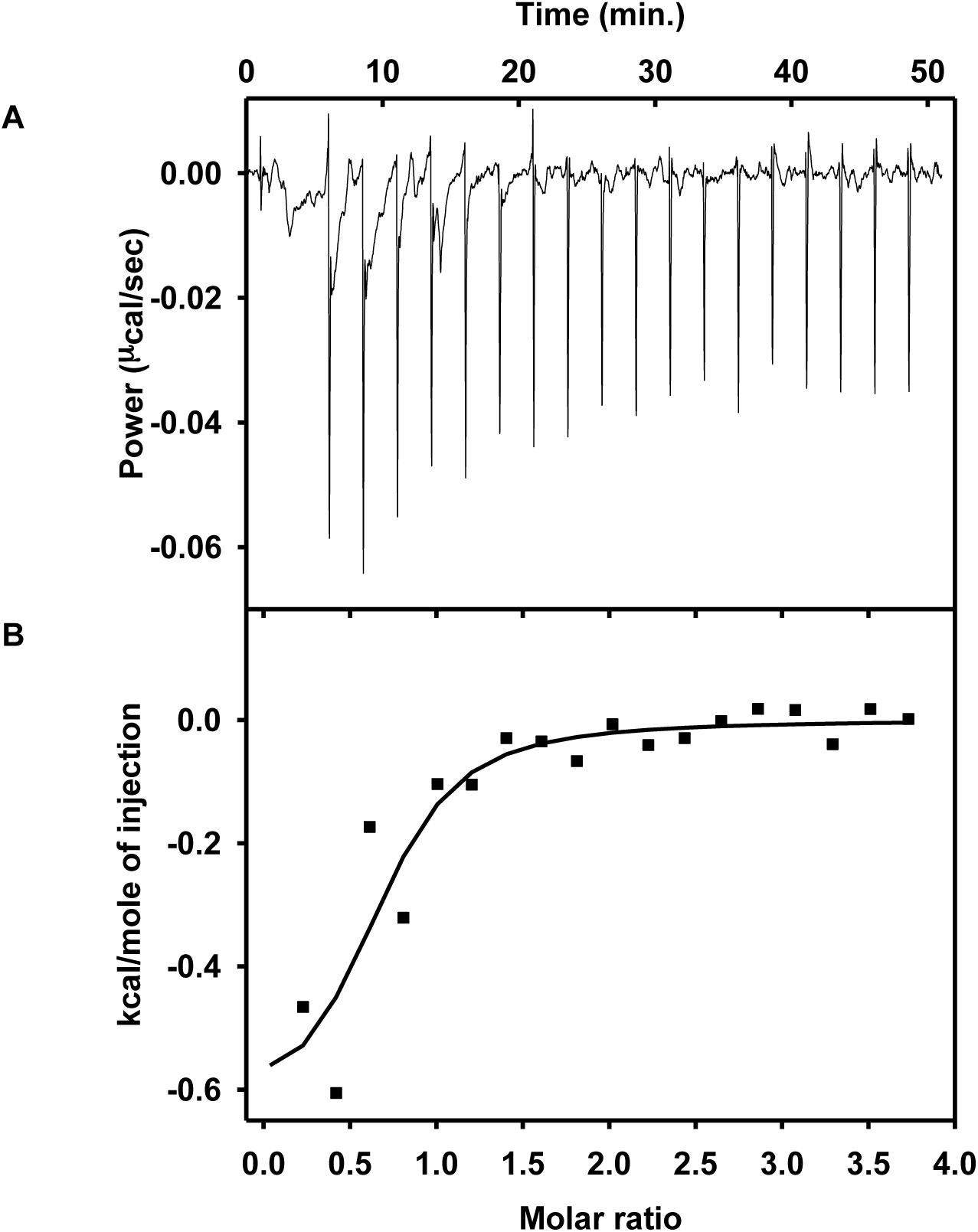
Interaction of Tau with CBMCs by ITC. (a) The differential power in terms of μcal/sec was plotted for each injections of L2. (b) The raw data was integrated as kcal/mole of injection versus molar ration of ligand, L2. The fitting of integrated points using one set of sites revealed the interaction of L2 with Tau with K_D_ of 5.09 ± 5.84 μM. The N value of 0.643 ± 0.132 suggests binding of one L2 to two Tau molecules.

### Cobalt inhibits the human Tau aggregation mediated cytotoxicity

Metals are an essential components of the brain at physiological levels but, their accumulation at higher concentration would be detrimental. Metals are known to exert toxicity in either of the following ways, by inhibiting PP2A or activating kinases such as cdk5/p25, GSK3β *etc*. The neurons isolated from hippocampus were treated with iron, this led to oxidative stress which ultimately caused Tau hyperphosphorylation *via*., cdk5/p25 complex.^50^ Similarly, studies in the rat primary cortical neurons showed that zinc caused phosphorylation of Tau at Ser262, which was marked as an early pathological events in AD.^51^ Here, the toxicity of these compounds were analysed by treating the SH-SY5Y cells with CBMCs in presence of Tau aggregates (Fig. 6E). The neuronal SH-SY5Y cells treated with Tau aggregates showed 59% cell toxicity at the concentration of 5 μM. However, this toxicity was arrested in the presence of CBMCs treatment. Interestingly, CBMCs displayed a dose-dependent inhibition of the cytotoxic effect induced by full-length Tau aggregates (Fig. 6F). Further, at CBMCs concentration of 100 μg/mL the cell viability was almost restored to 80%. These observations drive towards the conclusion that CBMCs are non-toxic to cells and moreover they reduce the toxicity of Tau aggregates, thus maintaining the cell viability.

**Figure 6.**
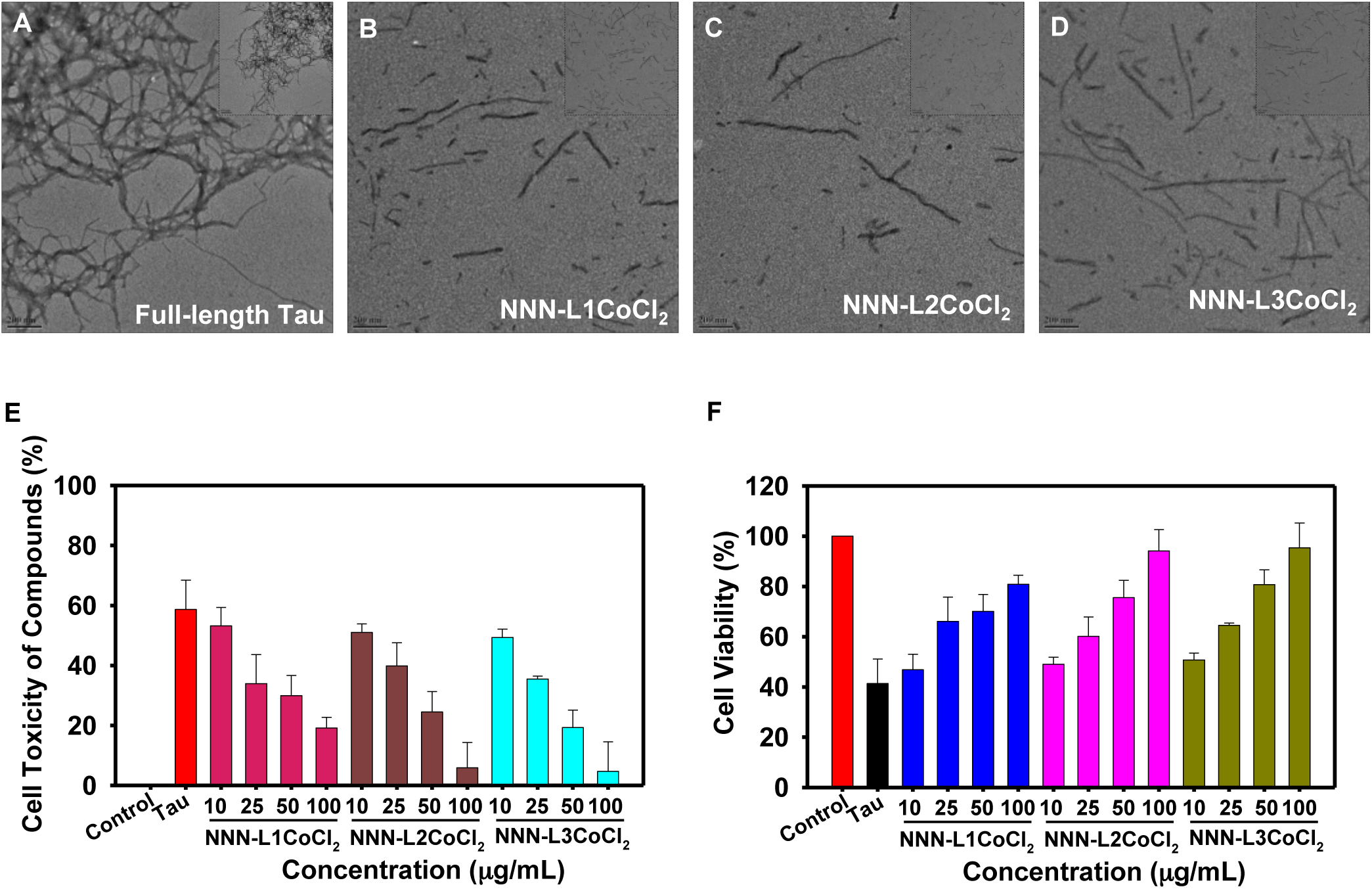
Disaggregation of Tau PHFs by CBMCs. (a) Aggregates of full-length Tau was observed by TEM after 120 hours of incubation in absence of metal complexes. (b, c, d) Upon prolonged incubation with L1, L2 and L3 the aggregates were destructed into shorter filaments. The inserts in each micrograph represent the morphology of disaggregation of Tau at magnification of 0.5 μm. (e) Cytotoxicity assay. The MTT assay signified that there was no toxicity of CBMCs on SH-SY5Y cells. (f) The CBMCs showed protective effect and helped in overcoming the toxicity of Tau aggregates at higher concentrations.

Further, at CBMCs concentration of 100 μg/mL the cell viability was almost restored to 80%. These observations drive towards the conclusion that CBMCs are non-toxic to cells and moreover they reduce the toxicity of Tau aggregates, thus maintaining the cell viability.

In summary, a newly developed molecular Co(II)-complexes (CBMCs) showed significant role in inhibition and disaggregation of Tau. The cytotoxicity studies on human neuroblastoma SH-SY5Y cells revealed that the present CBMCs are non-toxic in nature under *in vitro* conditions. We firmly believe that the present molecular cobalt(II)-complexes can be a potential therapeutic agent for Alzheimer’s Disease and related neurodegenerative diseases.

## METHODS

### Materials or Chemicals

MES, heparin, BES, BSA, BCA and ThS were purchased from Sigma. IPTG and DTT were purchased from Calbiochem. Other chemicals such as ampicillin, NaCl, EGTA *etc*. were purchased from MP. Cell culture related chemicals and plastics were purchased by Sigma and Thermo scientific Pvt Ltd.

Experiments were carried out using standard Schlenk techniques. All solvents were reagent grade or better. Deuterated solvents were used as received. Acetonitrile was refluxed over P_2_O_5_ and freshly distilled under argon atmosphere. Metal complexes (CoCl_2_) and other chemicals used in the reactions were used without additional purification. Thin layer chromatography (TLC) was performed using silica gel pre-coated glass plates, which were visualized with UV light at 254 nm or under iodine. The column chromatography was performed with SiO_2_ (Silicycle Siliaflash F60 (230-400 mesh). ^1^H NMR (400 or 500 MHz), ^13^C{^1^H} NMR (100 MHz) spectra were recorded on the NMR spectrometer. Deuterated chloroform was used as the solvent, and chemical shift values (δ) are reported in parts per million relative to the residual signals of this solvent [δ 7.26 for ^1^H (chloroform-d), δ 77.2 for ^13^C{^1^H} (chloroform-d). Abbreviations used in the NMR follow-up experiments: br, broad; s, singlet; d, doublet; t, triplet; q, quartet; m, multiplet. Mass spectra were obtained on a GCMS-QP 5000 instruments with ionization voltages of 70 eV. High-resolution mass spectra (HRMS) were obtained on a High-resolution mass spectra (HRMS) were obtained by fast atom bombardment (FAB) using a double focusing magnetic sector mass spectrometer and electron impact (EI) ionization technique (magnetic sector-electric sector double focusing mass analyzer).

**Scheme 1.**
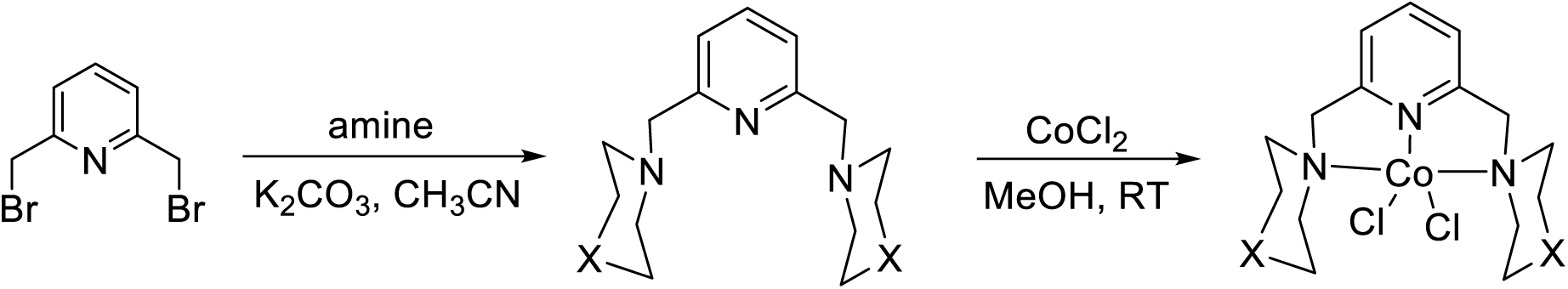
Synthesis of cobalt complexes (L1-L3)

### Synthesis of Ligands

#### 2,6-Bis(4-methylpiperazine-1-yl-methyl)pyridine (NNN-L1)

A solution of 2,6-bis(bromomethyl)pyridine (0.8 g, 3.0 mmol) in acetonitrile (45 mL) was added drop wise to solution of 1-methylpiperazine (0.669 g, 6.0 mmol) and K_2_CO_3_ (1.249 g, 9.0 mmol) in CH_3_CN (20 mL), the resulting reaction mixture was allowed to stir for 14 hours at 80-85°C, then cooled to room temperature, subsequently the reaction mixture was extracted with chloroform and water. The organic layer was collected, and evaporated in vacuum under the reduced pressure afforded yellow oil. Yield (0.82 g; 89%). IR (KBr): ν = 2945(s), 2520(m), 2042(m), 1452(s), 1029(s), 651(m). ^1^H NMR (500MHz, CHLOROFORM-d) δ = 7.59 (s, 1H), 7.28 (s, 2H), 3.66 (s, 4H), 2.54 (s, 8H), 2.46 (s, 8H), 2.28 (s, 6 H). ^13^C NMR (126MHz, CHLOROFORM-d) δ = 158.0, 136.6, 121.3, 77.3, 76.7, 64.4, 55.1, 53.2, 46.1. HRMS (EI): *m*/*z*Calcd for C_17_H_30_N_5_: 304.2501; Found: 304.2496.

#### 2, 6-Bis(piperazin-1-yl-methyl)pyridine (NNN-L2)

##### Step-1: synthesis of ^boc^NNN

solution of 2, 6-bis(bromomethyl)pyridine (1 g, 3.77mmol) in acetonitrile (40 mL) was added drop wise to solution of 1-boc-peprazine (1.4037 g, 7.54 mmol) and K_2_CO_3_ (1.56 g, 1.13 mmol) in CH_3_CN (20 mL), the resulting reaction mixture was allowed to stir for 14 hours at 85°C after cooled to room temperature, the reaction mixture was extracted in chloroform. The organic fraction were combined and dried over anhydrous and evaporated in vacuum, afforded yellow oil. (Yield 0.8 g; 80%). IR (KBr): ν = 3383(w), 2843(m), 2077(m), 1639(s), 1431(m), 1273(m), 1014(m), 559(m) cm^−^1. HRMS (ESI): Calcd. For C_25_H_41_N_5_O_4_ [M+H]^+^ 475.62; found 476.3244.

##### Step-2: synthesis of NNN-L2

^boc^NNN (0.8g) was dissolved MeOH (15 mL) and followed by added 1N HCl (8 mL), then the mixture was allowed to stir for 6hrs at 60°C. After cooling the reaction mixture to room temperature, it was neutralized with aqueous solution of 5% NaHCO_3_, then the solvent was evaporated in vacuum. The resulting sticky product was further dissolved in ethanol and filtered. The filtrate was concentrated in vacuum, afforded yellow oil product. Yield (0.75 g, 75%). ^1^H NMR (300 MHz, CDCl3) δ 7.68-759(t. 1H), 7.45-7.29(d, 2H), 3.92-3.64(s, 4H), 3.15-2.72(m, 8H), 2.71-2.42(s, 8H), 2.43-2.21(s, 2H).HRMS (ESI): calcd. For C_15_H_25_N_5_ [M+H]^+^ 275.39; found 276.21.

#### Synthesis of 2, 6-Bis(morpholinomethyl)pyridine(NNN-L3)

A solution of 2, 6-bis(bromomethyl)pyridine (0.3 g, 1.13 mmol) in acetonitrile (30 mL) was added drop wise to solution of morpholine (0.197g, 2.26 mmol) and K_2_CO_3_ (0.468 g, 3.39 mmol) in CH_3_CN (15 mL), the resulting reaction mixture was allowed to stir for 14 hours at 80-85°C, then cooled to room temperature, subsequently the reaction mixture was extracted with chloroform and water. The organic layer was collected, and evaporated in vacuum under the reduced pressure afforded colorless solid. Yield (0.282 g, 90%). IR (KBr): ν = 2800(m), 1575(m), 1454(m), 1298(m), 1111(s), 906(m). ^1^H NMR (500MHz, CHLOROFORM-d) δ = 7.65 - 7.52 (m, 1H), 7.31 (d, *J* = 7.6 Hz, 2H), 3.84 - 3.69 (m, 8H), 3.66 (s, 4H), 2.51 (s, 8H). ^13^C NMR (126MHz, CHLOROFORM-d) δ = 157.7, 136.7, 121.4, 77.3, 76.7, 66.9, 64.8, 53.7.

**Chart 1.**
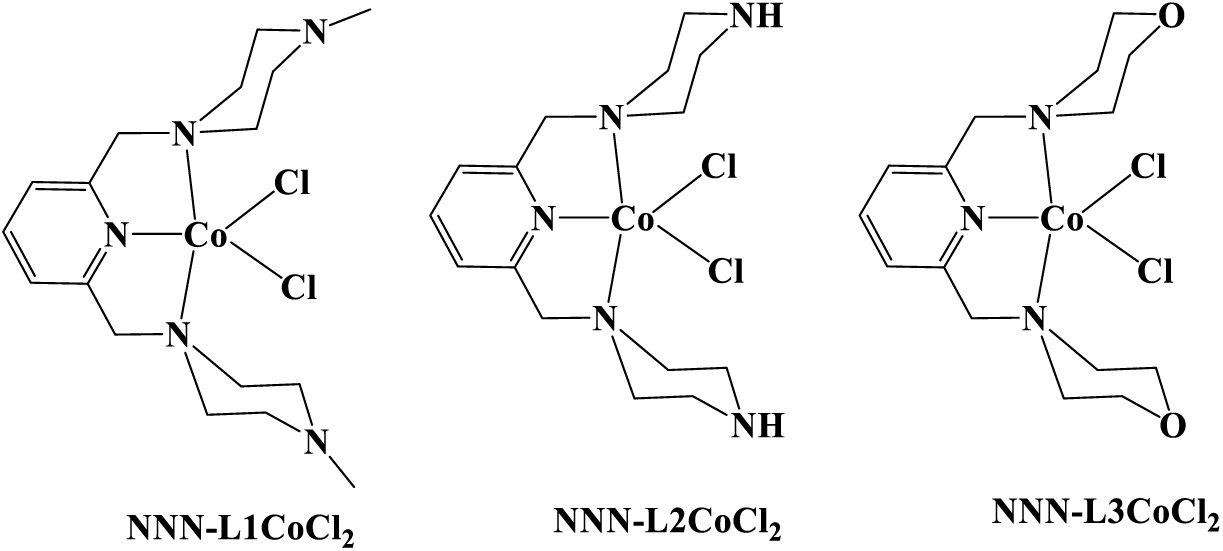
NNN-pincer type cobalt complexes

#### Synthesis of (NNN-L1)CoCl_2_

Cobalt chloride hexahydrate (0.312 g, 1.34 mmol) in methanol (15 mL) was added drop wise to solution of NNN-L1 (0.408 g, 1.34 mmol) in MeOH (15 mL) with stirring. The resulting reaction mixture was allowed to stir for 3 hours at room temperature. The resulting solution was evaporated under vacuum afforded the blue color solid; the solid was washed with diethyl ether and dried at air. Yield (0.54 g; 93%). IR (KBr): ν = 2924(s), 2314(m), 1612(m), 1462(s), 1207(m), 972(m). HRMS (EI): *m*/*z*Calcd for C_17_H_30_N_5_Cl_2_Co: 433.1210; Found: 433.1205.

#### Synthesis of (NNN-L2)CoCl_2_

Cobalt chloride hexahydrate (0.086 g, 0.36 mmol) was added to solution of NNN-L1 (0.1 g, 0.36 mmol) in MeOH (10 mL) with stirring. The resulting reaction mixture was allowed to stir for 3 hours at room temperature. The resulting solution was concentrated in a vacuum afforded the blue color solid; the solid was washed with diethyl ether and dried at air. Yield (0.132 g; 90%). IR (KBr): ν = 3446(w), 1633(s), 1460(m), 1165(m), 989(m), 613(m). HRMS (ESI): calcd. For C_16_H_28_Cl_2_CoN_5_ [M+Na]^+^ 405.23; found 429.24.

#### Synthesis of (NNN-L3)CoCl_2_

Cobalt chloride hexahydrate (0.129 g, 0.54 mmol) in methanol (8 mL) was added drop wise to solution of NNN-L3 (0.151 g, 0.54 mmol) in MeOH (10 mL) with stirring. The resulting reaction mixture was allowed to stir for 3 hours at room temperature. The resulting solution was evaporated under vacuum afforded the blue colored solid and the solid was washed with diethyl ether and dried at air. Yield (0.21 g, 95%). IR (KBr): ν = 2958(s), 2841(m), 1610(s), 1450(m), 1290(m), 1111(s), 999(m), 869(s), 815(m). HRMS (EI): *m*/*z* Calcd for C_17_H_28_N_3_Cl_2_Co: 403.0992; Found: 403.0987.

#### Expression and Purification of Tau

The recombinant full-length and four repeat Tau were expressed in BL21^∗^ strain of *E.coli*. Cells were induced with 0.5 mM IPTG after the OD at A_600_ reached to 0.5 to 0.6. The cells were allowed to grow at 37°C post-induction and were harvested by pelleting at 4000 rpm, at 4°C for 10 minutes. The pellet obtained was resuspended in 50 mM MES buffer pH 6.8 containing 1 mM EGTA, 2 mM MgCl_2_, 5 mM DTT, 1 mM PMSF, protease inhibitor cocktail and was lysed by using constant cell disruption system. The purification was done as described previously with minor changes.^52^ The concentrated protein was aliquoted, snap freezed and stored in −80°C until further used. The concentration was estimated using bichinchoninic acid (BCA) method.

#### Preparation of Tau aggregates

Tau was induced to aggregates formation as described previously with minor modifications.^52^ In presence of heparin (17,500 Da) at the ratio of 4:1,Tau was polymerized in assembly buffer containing 20 mM BES, pH 7.4, 25 mM NaCl, 1 mM DTT, 0.01% NaN_3_ and protease inhibitor cocktails. The reaction mixture was incubated at 37°C and the aggregates formation was monitored by ThS fluorescence,^53,54^ SDS-PAGE and TEM at certain time intervals. Tau protein was allowed to assemble in presence and absence of compounds in increasing concentrations with constant Tau concentrations of 0.91 mg mL^−1^. The changes in the conformation of Tau protein was monitored by CD spectroscopy.

#### Disaggregation assay

The potency of the metal complexes for disaggregating the preformed Tau aggregates was analyzed. Soluble Tau was incubated at 37°C for PHF assembly. The formation of aggregates was analyzed by ThS fluorescence assay and SDS-PAGE. Thus, formed aggregates were diluted to 0.91 mg mL^−1^ final concentration of 20 mM BES buffer, pH 7.4 and further the mixture was incubated with increasing concentration of metal complex as discussed earlier.

#### Thioflavin S fluorescence assay

5 μl of reaction mixture was diluted with 8 μM ThS in 50 mM ammonium acetate, pH 7.0 and added to 384 well plates in triplicates. Subsequently blank was also prepared for subtracting background fluorescence. The plate was incubated for 20 minutes in dark before measuring ThS fluorescence, at an emission wavelength of 521 nm by exciting it at 440 nm in Tecan Infinite 200 PRO multimode microplate reader.

#### SDS-PAGE analysis for Tau aggregates

The effect of the compounds on inhibiting the aggregates formation by Tau was observed by SDS-PAGE.^55,56^ The reaction mixture incubated with and without compound were collected at different time intervals of 0 hour, 24 hour and 72 hours (end point) and resolved 10% SDS-PAGE using miniVE Vertical Electrophoresis System from GE healthcare. Further the SDS-PAGE were quantified and analyzed using Gel Doc™ XR+ System and image lab software.

#### Soluble Tau Assay

The soluble Tau was studied in presence of metal complexes alone to analyze the conformational changes occurring due to the compound. 0.91 mg mL^−1^ was incubated for 1 hour at 37°C with and without different concentrations of 0.01 mg mL^−1^, 0.025 mg mL^−1^, 0.05 mg mL^−1^ and 0.1 mg mL^−1^ of metal complexes. At the end of one hour the samples were analyzed by SDS-PAGE, TEM and CD spectroscopy to monitor the formation of aggregates and change in Tau conformation, respectively.

#### CD spectroscopy

The conformational changes in Tau was analyzed by CD spectroscopy in the far UV region. Tau is a random coiled protein and upon aggregation it acquires β-sheet conformation. The impact of the compounds on preventing the formation of β-sheet structure was studied by CD spectroscopy. The spectra was collected as described previously, in Jasco J-815 spectrometer, by using cuvette with 1 mm path length.^57^ The measurements were performed in the range of 250 nm to 190 nm, with a data pitch of 1.0 nm, scanning speed of 100 nm/min. All the spectra were obtained at 25°C. The reaction mixture was diluted to 0.13 mg mL^−1^ in 50 mM phosphate buffer, pH 6.8. The effect of compound on soluble Tau was also studied by incubating Tau along with compounds alone at 37°C and the spectra was read at 25°C.

#### Transmission Electron Microscopy (TEM)

The extent of aggregates formed in presence of the metal complexes was analyzed by TEM (Tecnai T-20). The assay mixture was diluted to 0.04 mg mL^−1^ final concentration, spotted on the carbon coated copper grid of 400 mesh and incubated for 45 seconds. The excess Tau aggregates were removed by incubating the grid in water for 30 seconds and this was repeated twice. The grid was further stained by 2% uranyl acetate for 1 minute to observe the morphology of aggregates under TEM.

#### Size-exclusion chromatography (SEC)

The HMW species formed by Tau polymerization was analyzed by SEC.^58-60^ Tau protein was diluted to a concentration of 4.58 mg mL^−1^ in assembly buffer along with heparin in a ratio of 4:1 and incubated at 37°C in presence and absence of 0.1 mg mL^−1^ of NNN-L2CoCl_2_. Tau was subjected to SEC using superdex 75 PG in order to resolve aggregated Tau from the soluble, which was accessed as decrease in retention volume at different time points of 0, 2 and 24 hours in presence and absence of NNN-L2CoCl_2_.

#### Isothermal titration calorimetry

Isothermal titration calorimetry (ITC) was carried out to understand the thermodynamics behind Tau interaction with CBMCs. Here, the titration was done using 2.3 mg/mL of full-length Tau and 0.407 mg/mL of L2. The titrations were recorded in MicroCal PEAQ-ITC at 25°C. The titration was conducted by giving 19 injections, first injection of 0.4 μL was followed by injections of 2 μL each with 180 seconds spacing at stirring speed of 650 rpm. Tau and L2 were prepared in 20 mM BES containing 50 mM NaCl at pH 7.4. The samples were re-buffered, filtered and loaded. The sample cell was loaded with 200 μL of full-length Tau and syringe with 40 μL of L2. The data was analyzed in MicroCal PEAQ-ITC analysis software and fitted to one set of site model.

#### Cytotoxicity assay

SH-SY5Y cells were cultured in Dulbecco’s Modified Eagle Medium (DMEM)-F12 media (Gibco) supplemented with 20% FBS, 100 U/ml penicillin and 100 U/ml streptomycin. For the cell toxicity studies, sub-confluent cells were harvested by trypsinization and 25,000 cells/well were seeded in 96 well plate (100 μl/well). The cells were then incubated overnight at 37°C. Post incubation, the cells were treated with 100 μl of DMEM containing 5 μM full-length Tau aggregates, followed by the indicated amounts of the CBMCs. The full-length Tau aggregates alone was used as control. After 12 hours of incubation at 37°C, cell viability was evaluated using thiazolyl-blue-tetrazolium-bromide (MTT) assay. Each treatment was performed in triplicates. Briefly, 10 μl of 5 mg mL^−1^ MTT was added into each well and further incubated for 4 hours at 37°C. Later, 100 μl of DMSO was added into each well and the colour intensity was measured using ELISA Reader at 570 nm. The percentage of cell viability was calculated as following. Cells viability (%) = O.D. (570 nm) in the presence of full-length Tau with or without inhibitor^∗^ 100.

## ASSOCIATED CONTENT

### Supporting Information

Detailed Materials and Methods; Supplemental Figures regarding aggregation assay, soluble Tau and Size-exclusion data analysis, and electron microscopy data. This material is available free of charge via the Internet at http://pubs.acs.org.

## AUTHOR INFORMATION

**Corresponding Author**

To whom correspondence should be addressed:

**Dr. Subashchandrabose Chinnathambi,**

For synthetic chemistry part,

**Dr. Ekambaram Balaraman,**

## Notes

The authors declare no competing financial interest.

## Abbreviations used

AD: Alzheimer’s Disease;
Aβ: Amyloid β;
CBMCs: Cobalt-Based Metal Complexes;
NFTs: Neurofibrillary Tangles;
MTs: Microtubules;
PHFs: Paired Helical Filaments;
APP: Amyloid Precursor Protein

## Acknowledgements

This research was supported by the DST-SERB (SB/YS/LS-355/2013) and CSIR-NCL (MLP029526). EB acknowledges funding from SERB (SB/FT/CS-065/2013). Tau constructs were kindly gifted by Prof. Roland Brandt from University of Osnabruck, Germany and Prof. Jeff Kuret from Ohio State University College of Medicine, USA. NVG acknowledges to UGC and SPM, and VGL thanks to CSIR for fellowships.

## Table of contents

Tau attains pathological conformation during Alzheimer’s disease, which ultimately leads to neurodegeneration. The synthetic rationally designed molecular cobalt complexes (CBMCs) prevents the polymerization of Tau protein, as well as destabilizes the pre-formed fibrils. Thus, CBMCs are identified as a hit compound with significance to prevent AD as evidenced by various biochemical and biophysical analysis.

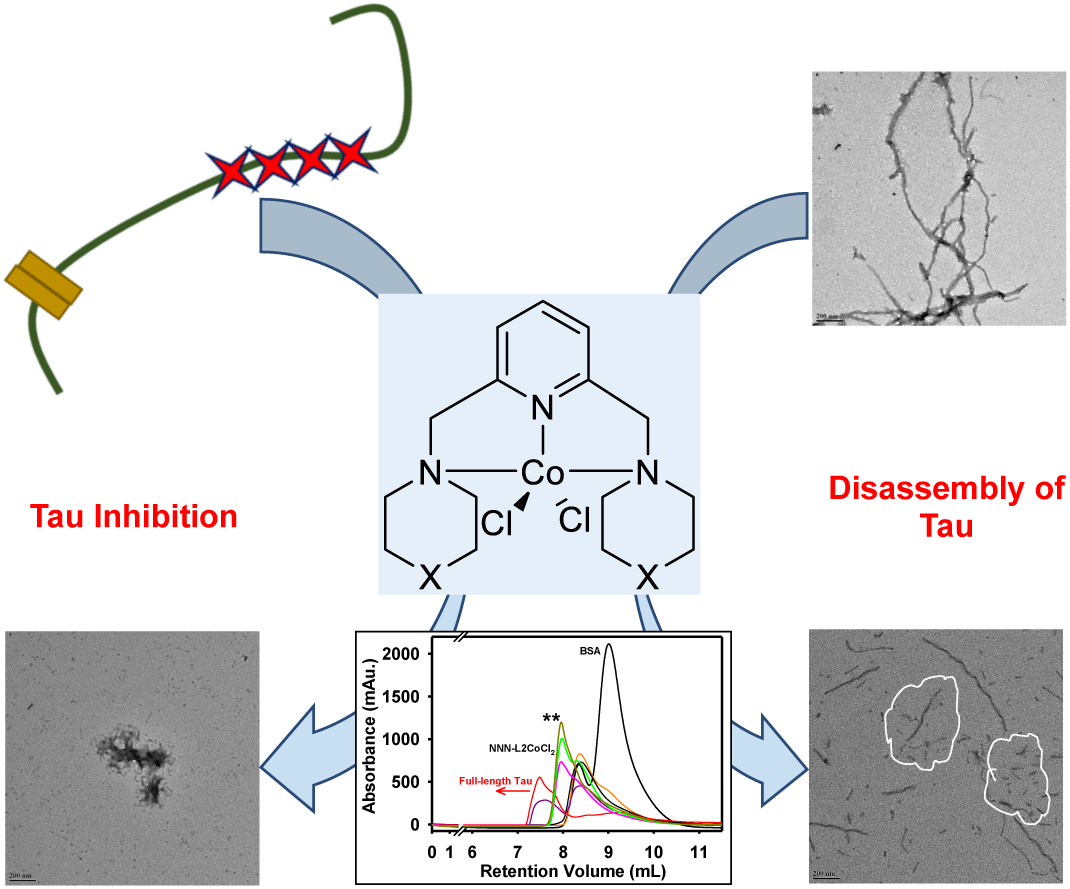

